# A workflow for microclimate sensor networks: integrating geographic tools, statistics, and local knowledge

**DOI:** 10.1101/2024.09.13.612939

**Authors:** David H. Klinges, Jonas J. Lembrechts, Stijn Van de Vondel, Eric Greenlee, Kian Hayles-Cotton, Rebecca A. Senior

## Abstract

Wireless environmental sensors have become integral tools in environmental and conservation research, offering diverse data streams that complement traditional inventory-based surveys. Despite advancements in sensor technology, the ad-hoc nature of site selection for sensor deployment often limits the potential of collected data. In this paper, we argue for the importance of informed site selection to capture environmental variation effectively. We introduce a comprehensive step-by-step practical guide for environmental sensor site selection and network deployment, drawing on experiences from diverse geographic locations and focusing on microclimate monitoring as a representative environmental variable. The workflow integrates Geographic Information Systems (GIS) tools, local community-based knowledge, and statistical methods to provide adaptive and iterative guidelines for both new and expanded sensor deployments. We demonstrate the workflow’s applicability across three distinct settings: arid montane deserts in Oman, urban and rural gardens in Belgium, and humid forested landscapes in Madagascar. To facilitate the workflow’s implementation and reproducibility worldwide, we provide a modular software supplement with flexible user input for robust, data-driven and interactive site selection. Critically, our workflow underscores the importance of equitable collaboration with local stakeholders, addresses challenges in sensor deployment, and offers a practical tool to enhance the effectiveness and efficiency of environmental sensing across disciplines including ecology, meteorology, agriculture, and landscape design.

## Introduction

In recent decades, wireless environmental sensors – any device that can automatically collect and store information on its surroundings without an external power source – have become a ubiquitous tool in environmental and conservation work (Bush *et al*. 2017). Camera traps, soil chemistry kits, microclimate loggers, and other technologies provide a diversity of data streams that can supplement inventory-based surveys. Decreasing costs, improved battery power, and enhanced portability have also made deploying sensors in remote locations increasingly feasible, to the benefit of many practitioners (Buratti *et al*. 2009). Given that much ecological research is primarily focused on particular biological processes or chosen taxa, the designation of sites for wireless sensing is usually ad- or post-hoc: sites are chosen based on ease of access, “habitat quality” for focal organism(s), and feasibility for biological surveys (Carvalho *et al*. 2016; Nuñez-Penichet *et al*. 2022), with sensors then deployed at those sites to capture conditions that are most proximal to focal organisms or habitats.

Ad-hoc sensor measurements may be useful to understand conditions at a given spatial point and to answer initial research questions related to organismal ecology or biodiversity. Yet informal approaches to selecting sensor sites, such as “tacking on” sensors to locations primarily chosen for taxonomic surveys, have consequences that limit the potential of sensor data. First, the drivers of biodiversity in space or time (which may have justified site selection) are not always the same drivers of environmental patterns that a sensor attempts to measure (Abrahms *et al*. 2020). Conventional biological sampling, therefore, may not effectively capture the full range of environmental variation. Environmental sensor networks and corresponding datasets can also have new life cycles beyond a single study, and informed site selection is key for making data useful for downstream applications. For example, the proliferation of landscape statistics, satellite imagery, and corresponding software for merging spatial models with in-situ measurements (Bush *et al*. 2017) has made it increasingly feasible to leverage sensor measurements for spatial prediction across a region (Haesen *et al*. 2023) or for forecasting future conditions (Thomas *et al*. 2023). Additionally, sensors or their measurements can be incorporated into regional, national, or global networks to synthesise insights across scales (Lembrechts *et al*. 2020, 2021). Yet if sensors do not adequately sample a region’s heterogeneity, their direct measurements – and especially any extrapolated predictions such as forecasts – may lead to poorly informed insights or management decisions (Acevedo *et al*. 2015).

Furthermore, scientific questions, longevity of research insights, and utility of products for real-world applications all rely on equitable collaboration between local stakeholders and scientists or managers (who may be foreign to the region of interest). Given that sensors may be deployed for extended periods to provide data with broad utility, co-design of site selection with stakeholders is especially important. Sensors may be aesthetically displeasing or arouse suspicion in natural settings, may be at risk of theft (Farris *et al*. 2017), or can even disrupt use of an area by humans or other species (Carroll *et al*. 2020), all of which provide rationale to not only request permission from local communities, but also understand their values.

Since environmental sensors deployed by ecologists are usually secondary to biological variables, important considerations such as the necessary sensor sample size, deployment locations, and best practices to address sensor failure have received little attention in the ecological literature. Protocols are dependent on research questions, sensor types, and budgetary and temporal constraints, yet there are general principles that unify decisions across contexts. Statistical power analyses, often forgotten after graduate training, remain useful for estimating necessary sample sizes to draw inference on a pattern or process (McDonald 2014). Randomised or stratified site selection (Carvalho *et al*. 2016; Gillespie *et al*. 2017; Nuñez-Penichet *et al*. 2022) can leverage spatially gridded remote sensing to understand environmental heterogeneity across a landscape (Cordell *et al*. 2017; He *et al*. 2015), leading to more balanced and scientifically representative sampling (Chadwick *et al*. 2020; Lembrechts *et al*. 2021). Given that most sensor deployments are tailored for particular projects or landscapes, insights on best practices are rarely published or shared.

To unify efforts, we provide a practical and generalised workflow for site selection for environmental sensors (Figure 1). We constructed this workflow drawing upon our broad experiences in microclimate-centred monitoring from across six continents. Here, we define microclimate as the local climate conditions within one metre of the earth’s surface, and/or below vegetation, that organisms and ecosystems are exposed to (Geiger 1942; Kemppinen *et al*. 2024). Microclimate serves as an excellent example of a suite of environmental variables that are broadly relevant for ecology and society, and for which both remote sensing and *in-situ* insights are tremendously valuable (Zellweger *et al*. 2019). Our workflow leverages GIS tools and data while also incorporating key insights on landscape configuration, site access, and design feasibility that can only be learned locally. Such local knowledge is especially critical where satellite imagery is less reliable, such as cloudy areas or those at high latitudes, and is integral to applying ecological insights for policy and management. We provide an iterative guidelines that are flexible in the face of unforeseen events, especially for remote fieldwork for which frequent visitation is challenging. We describe the workflow in nine discrete steps and demonstrate its application in three disparate settings: urban gardens in Belgium, arid and sparsely-populated Oman, and humid forest and croplands in Madagascar (Figure 2). Importantly, we provide a modular, well-documented code supplement, which generates coordinates for recommended sensor deployment within a user’s region of interest based upon their budget, spatial remote sensing grids, and other user inputs, to make our workflow executable and reproducible across the planet. We also discuss common challenges, practices to incorporate local knowledge, and the prospects of assimilating our workflow with innovative tools such as Internet of Things (IoT) sensing. The workflow and corresponding insights developed here have broad utility for environmental sensing by academic, governmental, and local community stakeholders to improve applied science, reduce costs, and make fieldwork more productive.

**Figure 1.**
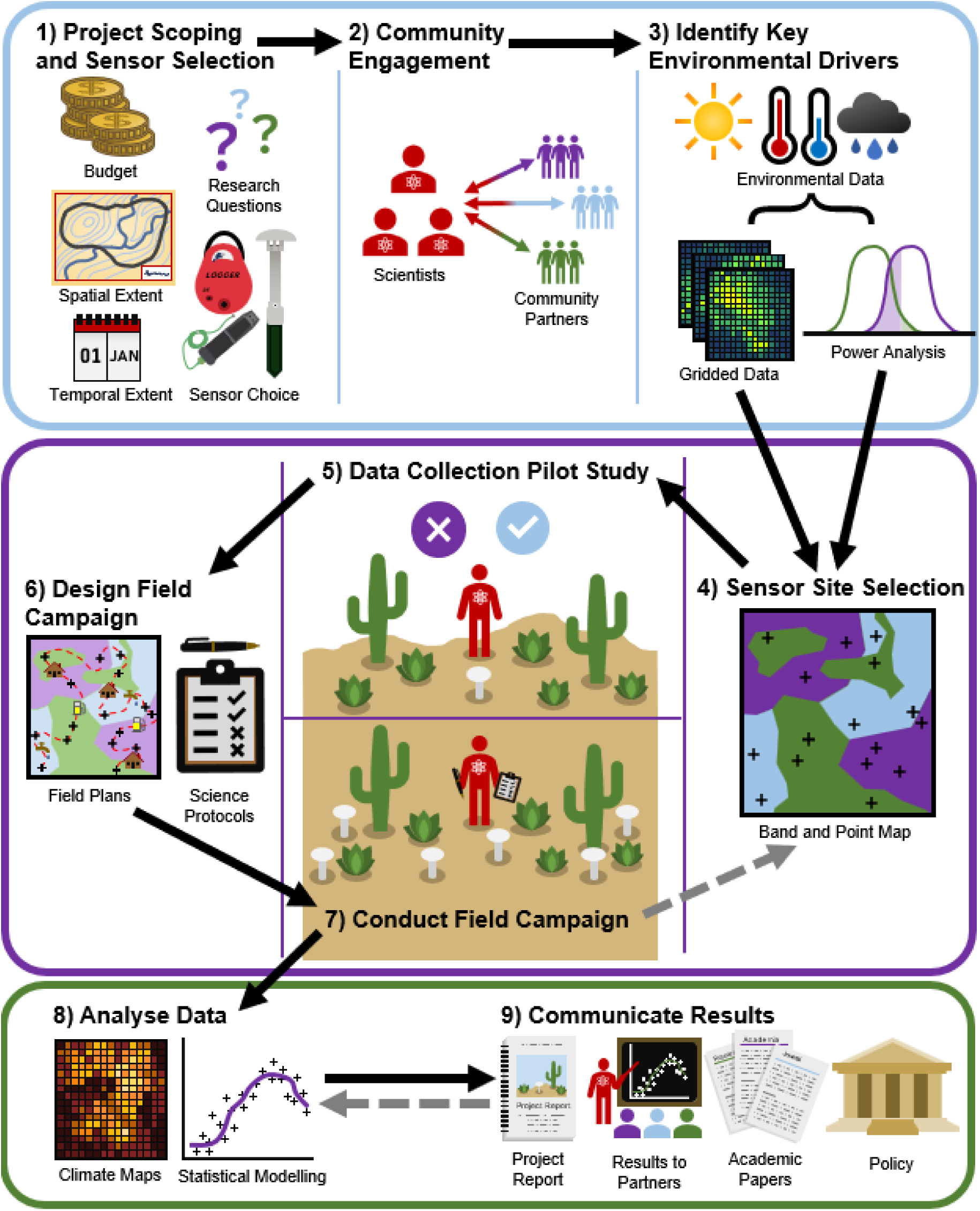
Visual depiction of our nine-step workflow, grouped in three sections. Project scoping (1), community engagement (2) and identification of key environmental drivers (3) should lay the groundwork before the field campaign, while site selection (4), pilot study (5), and design (6) and execution (7) of the field campaign form the core of the microclimate monitoring study. After the field campaign, data needs to be analysed (8) and results communicated (9). Arrows indicate the order of steps, and occasionally include iterating through earlier steps (grey dashed arrows).

**Figure 2.**
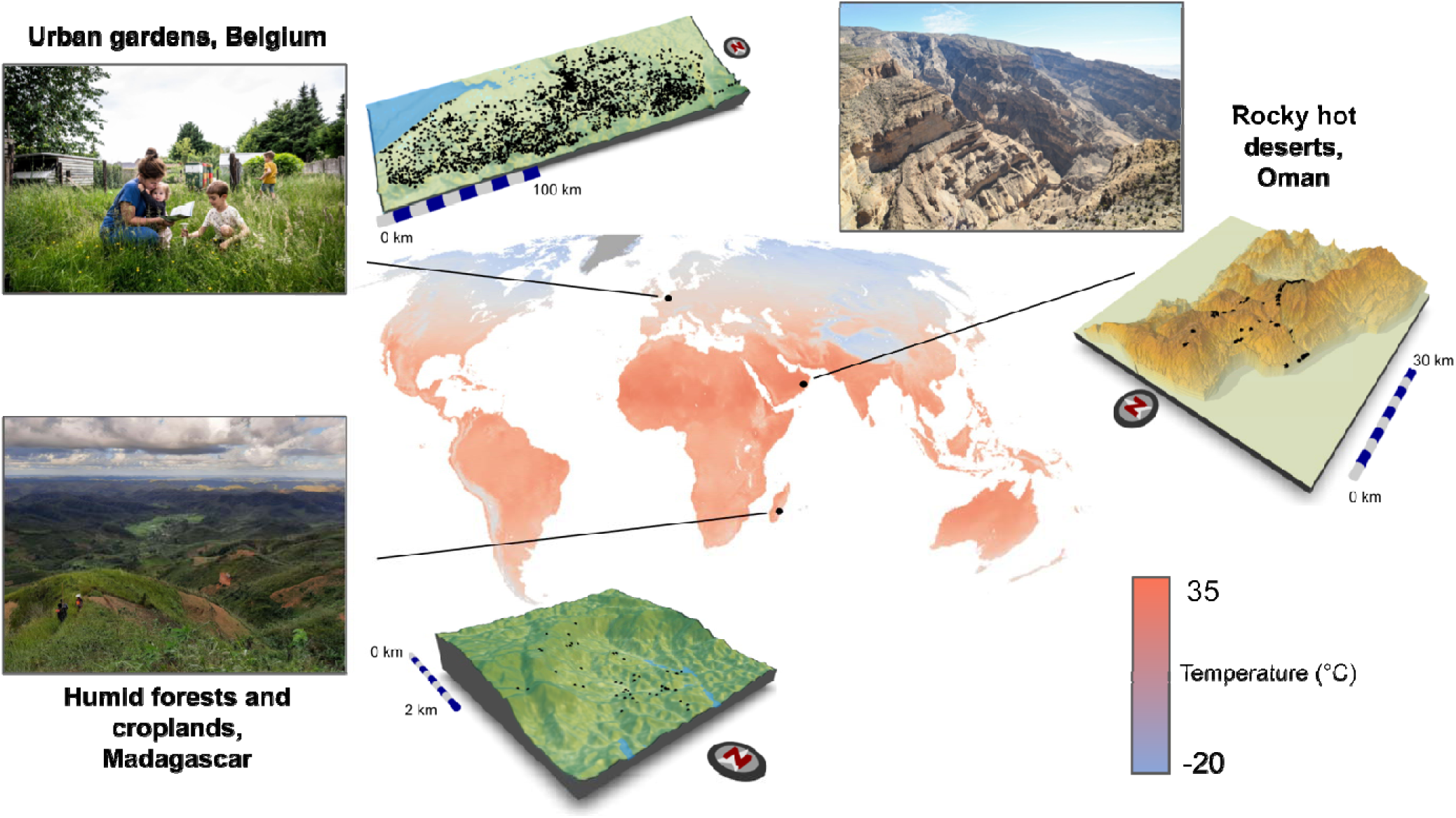
The three case study regions in which the sensor network implementation workflow was applied. Insets depict each landscape and a topographical model of each region, with black points indicating where microclimate sensors were deployed. Background global map of mean annual soil temperature from Lembrechts *et al*. (2022).

### Step-by-Step Workflow

We provide our workflow in nine steps that standardise study design, data collection, and result communication for research using microclimate sensor networks (Figure 1).

While previous work has provided guidance on developing countrywide networks (Lembrechts *et al*. 2021), our workflow is geared towards sensor deployment across landscapes (areas < 100 km^2^). Our workflow approaches microclimate network design from a practical perspective by summarising all steps needed from design to implementation, while confronting site access and feasibility issues. Protocols outlined here focus on decision-making related to sensor deployment locations at the plot level (e.g. within a 10m-30m footprint). This target resolution matches most fine-grain satellite imagery, and is typically sufficient for capturing geographical heterogeneity and differences in vegetation types (e.g., forest versus grassland). Precise sensor deployment within the 10m-30m spatial area can then be defined based on context and goals.

#### Step 1: Project Scoping and Sensor selection

The first step is to decide the scope of the project, which encompasses research question(s), budget, spatial extent, temporal duration, and potential stakeholders. Projects may have these components previously determined due to funding structures, existing partnerships, and timelines. For example, a sensor deployment may be part of a broader biological and environmental monitoring agenda with a fixed budget for sensors. Project goals, budget, community contacts, and logistical travel constraints thus determine the spatial extent and temporal duration. Alternatively, if a sensor network is the core focus of a project, the spatial scope and sensor data types (e.g., temperature vs. water chemistry) will inform a necessary budget and identify community stakeholders. Regardless, we emphasise that scientists iteratively fine-tune goals and project scope throughout this workflow.

For microclimate sensor deployment, delineating research questions entails identifying both the microclimate response variables of interest (e.g. temperature, humidity, light availability, soil moisture) and the potential sensors to measure them. Which sensor to use is one of the most critical decisions to take when establishing an environmental monitoring network. While the optimal sensor will differ depending on research questions, study system and budget, we summarise a few general considerations in Box 1.

Clarifying the scope is important from the beginning, but not every component needs to be fully planned at this stage. For example, although spatial extent and temporal duration are important to determine early, the spatial and temporal resolution can be decided later (Steps 4 and 5).

###### BOX 1: General considerations for selecting microclimate sensors

- *Accuracy*: many cost-effective sensors suffer from low accuracy, especially when measuring near-surface air temperature in open terrain (Maclean et al. 2021). Most thermometers absorb radiation, thereby introducing considerable bias in temperature measurements, which has led to the widespread practice of shielding thermometers. However, the presence of a shield may prevent measurement of the microclimates of interest (Terando *et al*. 2017). Ultrafine-wire thermocouple sensors avoid the need for shielding, with many consumer-grade options (see Maclean *et al*. 2021 for a thorough treatment of near-surface air temperature measurement for ecological applications). Accuracy is usually less of an issue in soil or shade.
- *Ample memory space*: sensors with sufficient memory capacity ensure data integrity over long periods, allowing finer temporal intervals, fewer field visits, and mitigating the risk of data loss.
- *Multi-sensor functionality*: sensors measuring multiple parameters simultaneously provide more comprehensive insights into microclimate dynamics, though can be more expensive or compromise on quality.
- *Long battery life*: extended battery life minimises the need for frequent replacements, ensuring uninterrupted monitoring.
- *Durability*: robust sensors or casing can withstand harsh environmental conditions to ensure reliable long-term monitoring.
- *Standardisation*: standardised protocols simplify data collection and analysis, facilitating comparability across studies. This includes both hardware (e.g. battery size) and software; for the latter, we advise use of brands that provide data in comma-separated format (CSV) with freely available (and, ideally, open-source) programs for processing.
- *Installation restrictions*: sensor selection may be informed by sensor size (e.g., to avoid influencing the study organisms), sensor visibility (e.g., to avoid theft or vandalism), sensor height (e.g., no aboveground part to avoid issues with agricultural management) or sensor depth (e.g., project requires monitoring below topsoil).
- *Shipping restrictions*: regulations governing the transportation of sensors and batteries may pose logistical challenges.

#### Step 2: Community Engagement

Effective microclimate monitoring often relies on accessing diverse microhabitat(s) across a landscape, for which it is important to identify and consult local landowners and stakeholders from the outset. By building partnerships with groups who possess local knowledge, the collected microclimate data can better represent the study area and increase the project’s impact and longevity. Creating a direct route from data to stakeholders also builds buy-in among community members who likely have questions of their own about the landscape. By better understanding the direct benefits of microclimate research, communities are more likely to provide endorsement or project support.

Relevant communities to involve in a microclimate measurement campaign will vary across projects, but guiding principles to consider are goal alignment, data re-use, and respect for local knowledge (Rothrock *et al*. 2023). To recruit diverse partners with capacity for taking on projects, Leone *et al*. (2023) recommend hosting a multi-round “Request for Partnership” call among candidate groups. When a project requires collecting data on or travelling through private land, permissions should be acquired as early as possible. Even in cases where land is not legally private, community members may attach special value to these locations. Community conversations may also foster engagement, such as citizen science (e.g. garden owners in the Flemish case study below). Partnerships will likely include both formal components laid out with official organisations and informal components built naturally during time spent in community spaces.

Genuine stakeholder engagement often requires flexible timeframes and incentive structures. When approaching communities, leverage past connections to build trust. Explain the proposed methods and purpose of the project and listen to any suggestions to better align the project with community goals. It is important at this stage to remain realistic about the project’s impact and not overpromise on management or policy ramifications. When situationally appropriate and funding is available, we recommend transparent and user-friendly access to data (summaries), and/or financially compensating community members for their time spent planning or monitoring site deployments. Some groups may be happy to volunteer their time, but others will be inconvenienced by contributing towards a project that they may never benefit from.

We cannot be exhaustive on the complex but important topic of community-engaged research, and instead refer the reader to academic literature on the many benefits and pitfalls of this topic (Douglass *et al*. 2019; Norström *et al*. 2020; Sandbrook *et al*. 2023). We offer further insight in the “Co-development across the Workflow with Stakeholders” section below.

#### Step 3: Identify Key Environmental Drivers

After project aims have been delineated and relevant partners have been engaged, one should determine the important drivers of the microclimatic variables of interest. Spatial data on these drivers will serve as input for the location selection algorithm (Step 4). Many foundational texts have identified and reviewed the most important drivers of microclimate (Bramer *et al*. 2018; Geiger 1942, De Frenne et al. 2021), which we will only briefly summarise here.

Broadly speaking, microclimate is influenced by local-scale variation in vegetation, terrain, and soil conditions (Geiger 1942), which should be captured through appropriately chosen sensor locations. Importantly, drivers of microclimate will not play equal roles across all ecosystems and applications. Furthermore, interactions between drivers are often landscape-specific. For instance, the importance of vegetation cover depends on the topographic complexity of the region. Our code supplement automatically downloads and processes public spatial data that provide measurements or predictions for many relevant microclimate drivers: macroclimate, elevation, slope, aspect, soil temperatures, and categorical land cover. Our code supplement standardises projections, resolutions, and extents of these layers to suit the requests of users (e.g. based upon a user-specified coordinate system and target region).

Just as important as collating spatial data is identifying which drivers are not represented in public domain spatial gridded data. Local land management zoning, for example, may not be readily available online, and either may need to be manually digitised (e.g. from paper or PDF maps), collected during site visitations, or obtained from relevant stakeholders. Our code supplement allows users to input custom digitised maps to supplement default environmental layers. Certain spatial information may also be sensitive or personal (e.g. household locations or observations of species threatened by poaching), and care should be taken to maintain confidentiality where necessary. Given the many sources of relevant spatial information, we recommend prioritising a short list of potential drivers, for the sake of pragmatism, parsimony, and reproducibility. Our code supplement provides power analyses to aid decision making: calculating the number of sensors needed to effectively capture variation in a set number of microclimate drivers, or calculating the number of drivers for which variation can be explained by a set number of sensors that a budget can support. Thus, this step is completed once both sensor quantity and the set of chosen relevant drivers are established.

#### Step 4: Sensor Site Selection

Once the sensor count is defined, it is time to designate sensor locations. Focusing solely on physical space for site selection may not optimally capture the non-uniform distributions of microclimate within a landscape, hence we provide a more effective strategy that includes the important spatial layers identified in Step 3. This method integrates systematic multivariate sampling of environmental space of a chosen landscape. As implemented in our code supplement, we make use of ordination (principal component analysis for continuous input layers, or factor analysis for a mix of continuous and categorical input data; automatically determined from the user’s spatial layers) to select representative locations that cover the maximum range of microclimatic conditions. Through this multivariate analysis, our program creates a set of environmental ‘bins’ across multiple dimensions, grouping locations with similar environmental conditions (and therefore anticipating similar microclimate; Figures S1-S2). By evenly distributing sensors across these bins, sampling can maximise microclimate coverage with a limited number of sensors, and also reduces dependency on any given sensor (important if the likelihood of sensor failure is high). Our code supplement is highly flexible, allowing the user to designate only specific eligible locations (e.g. if just one habitat type is of interest), and with the option to incorporate pre-existing sites so that new selected sites are sampling within complementary environmental space. Our program also allows for an iterative approach of updating sensor locations if certain sites are deemed inaccessible or invalid (see Step 8).

The distance between chosen sites may be important for ecological or logistical reasons. In addition to selecting sites based upon environmental variation, the code supplement provides functionality to set a minimum distance between all sites (if spatially extensive sampling is desired) and to set a maximum distance between each site and the nearest site (i.e. to facilitate multi-site access).

For pre-defined budgets in which a per-sensor cost is provided, the analysis determines the maximum number of sensors, and consequently the environmental variation one can sample. Conversely, when budget flexibility exists, the program will suggest an ideal number of sensors.

#### Step 5: Data Collection Pilot Study

When establishing an entirely new sensor network, conducting a pilot study to verify that chosen sensors, accessory equipment, and deployment protocols are all functional and interoperable is recommended. We advise using such a pilot to identify the appropriate choice of sensor housing and shielding, which can have tremendous impacts on microclimate observations (Terando *et al*. 2017). Furthermore, testing deployment and data harvesting protocols can help calculate the time needed for field installation and reduce the risk of misconfigurations. A pilot study also tests if the sensor and its data follow the expectations shaped in Step 1.

Other considerations during a pilot include site access and sensor longevity in the field. For fieldwork in rugged terrain or requiring long distances hiked by foot, calculating total weight per site (sensor and all accessory equipment) can be useful for feasible deployment. For long-term deployment, water- and heat/cold-proofing is especially crucial, as proprietary casing of a sensor may not be sufficient to survive adverse weather events (Mickley *et al*. 2019), and both wild and domesticated animals can cause severe damage to sensors (Wild *et al*. 2019). Battery life and durability are critical concerns, as a dead battery can potentially corrupt prior data. If potentially restrictive, we recommend testing battery lifetimes (which may be less than marketed lifetimes) within simulated field conditions before deployment. Battery lifetime may be sensitive to the temporal interval of data collection, and therefore a pilot study and considerations about the minimum temporal resolution required for the question at hand can inform the ideal trade-off between temporal resolution and span of deployment before battery failure. We encourage temporal sampling of no coarser than every two hours, as microclimate relevant for most organisms varies at the hourly scale (Maclean *et al*. 2021). Measuring battery life can inform the timing of revisits to field sites, and some IoT sensor networks also can provide real-time metadata on battery status (Callebaut *et al*. 2021; and see “IoT as the Future of Sensor Networks” below).

#### Step 6: Design field campaign(s) and survey protocols

With sensor locations and backups established in Step 4, field campaigns for sensor installation, check-ups, data retrieval, and metadata collection can be designed. When local partners (e.g., site managers) or citizens are involved in sensor installation, researchers must take measures to facilitate this procedure. This includes providing appropriate installation instructions and/or information sessions, transporting sensors to local partners (which may include logistical challenges associated with delivery companies, transport of batteries, and import fees), and communicating timelines to all partners. When these decisions have been made, formalising a memorandum of understanding (MoU) summarising project goals, expectations, and protocols among partners ensures aligned expectations. This involves delineating financial responsibilities (e.g. who is in charge of sensor purchase and replacement, and potential stakeholder compensation?), work divisions (who is in charge of sensor installation, retrieval, data analysis?), and communication protocols (which outcomes will be communicated to whom, when, and by whom?). This pivotal phase marks the realignment of project scope, resources, and collaborative commitments in harmony with refined insights and goals.

#### Step 7: Conduct Field Campaign(s)

Of the many steps likely needed during a field campaign to fulfil one’s research goals, here we focus on those most pertinent to deploying sensors and updating site selection. Sufficient documentation of sensor locations should be made during the field campaign; coordinates estimated by a handheld GPS or smartphone are critical, yet serve as a bare minimum. Additional steps to aid sensor retrieval include written notes on sensor height/depth and orientation, as well as photographs of exact sensor location from different distances away and angles of approach. Depending on human and animal threats to sensors (e.g. theft, trampling, etc.), consider flagging sensors with visible ribbon, locking sensors to sturdy objects, or caging/camouflaging sensors for protection. Finally, it is wise to record the time when the sensor was turned on and fully deployed.

Almost inevitably, plans will require reconsideration when confronted with the realities of new sites and sensor deployments. We therefore encourage an iterative approach to sensor site selection, as implemented in our code supplement. This program updates sites of yet-to-be deployed sensors according to new information gathered on-the-fly during fieldwork. If a subset of algorithm-selected points are deemed inaccessible or inappropriate upon site visit, the user can provide a CSV to the program of which points remain eligible. The algorithm will then choose new points that complement the sampled environmental space of established sites. This procedure can be iterated until all points have been visited and confirmed as viable deployment locations.

This iterative approach requires access to a computer. Although we were able to implement such an approach with personal laptops in our case studies, two of which are off-grid and remote (Madagascar and Oman), we recognize that this may not be feasible or recommended for all projects. We have therefore designed the code supplement to be intuitive for a wide audience, so that the program could be run by off-site collaborators, requiring only a simple spreadsheet of eligible points for an additional selection iteration.

#### Step 8: Result documentation, analysis, and visualisation

Several methods and tools available using the R programming language (R Core Team 2024) may be useful for processing and analysing microclimate sensor time series. The package *myClim* (Man *et al*. 2023) can process observations from TOMST and Onset HOBO loggers, and routines can be easily customised to support Lascar, iButton, and Logtag measurements. *myClim* allows data cleaning, homogenising, and checking for potential errors. For constructing continuous thermal regimes from temporally sparse measurements, we highlight the *sinc* package (von Schmalensee 2023). The package *climwin* (Bailey & Pol 2016) can help identify the temporal windows (e.g. end of winter, or height of rainy season), for a given measured climate variable, that are the most important for explaining variation in a specified ecological response variable. Our code supplement automatically provides users with several sets of visuals that contextualise sensor deployment locations within spatial and environmental variation, including maps of sensor locations with different environmental conditions as base layers, histograms of sampled environmental variation overlaid on total environmental variation, and the areas of multivariate environmental space represented by chosen sites (Figures 3, S1-S4). These visuals aid communication of final sensor locations to project partners, which should be supplemented with written descriptions of locations.

**Figure 3.**
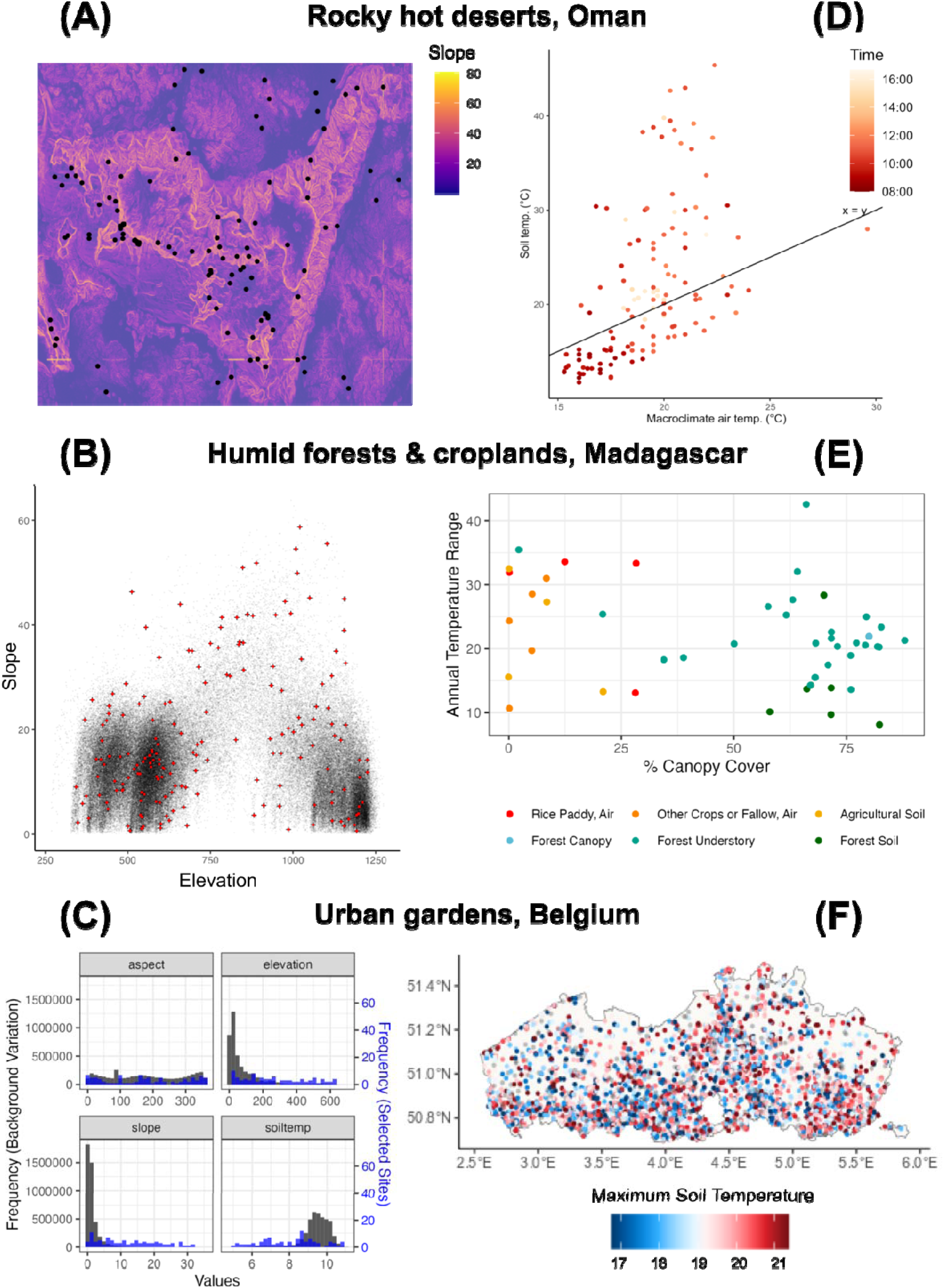
Data visualisations can both depict environmental representation of chosen sensor locations for internal review by project researchers (panels A - C) as well as communicate project results to target external stakeholders (panels D - F). Panels A - C depict figures automatically generated by the code supplement for a user’s requested landscape for any terrestrial location on earth, here displayed for each of the three case study regions: Belgium (top), Oman (middle), and Madagascar (bottom). The code supplement selects sensor locations that maximise representation of multivariate environmental variation, which is then displayed in several formats: (A) maps of sensor locations with each environmental layer in the background (here exemplifying slope across Oman); (B) scatterplots of chosen locations (red points) across existing environmental variability within the landscape (here plotting slope against elevation in Madagascar); (C) histograms of the distribution of selected sites across the distribution of each environmental variable across the landscape (here displaying variables for Belgium). After sensors have been deployed, project members should cater visuals to the target stakeholders for delivering results: (D) demonstrating how topsoil temperatures can decouple considerably from macroclimate depending on microtopography in the Western Hajar central massif of Oman; (E) depicting the range of microclimatic temperatures for different land use types for farmers in Madagascar; (F) maps of thermal summaries (here, hot extremes) in Belgium that were provided in a national newspaper to engage the public.

#### Step 9: Communicate results

While communication with various stakeholders and the broader scientific community will occur throughout a project, it is important to deliberately inform partners of the results at the completion of key milestones. Beyond interacting with the research community through scientific papers and conference talks, local stakeholders will likely warrant different forms of communication, and it is imperative to re-engage them even if their involvement may have ended. Integrating conversations into existing community forums or demonstration workshops can encourage participation and local buy-in, though care must be taken to repackage materials into accessible language and graphics to promote equitable outcomes (Figure 3; Douglass *et al*. 2019). Such gatherings can also provide opportunities to explore future projects together. Beyond in-person meetings, consider disseminating project results to local news outlets or making them freely available online (see Belgian case study below). Communication outside of typical academic routes can have great benefits for both the research team, who build closer ties with local partners, and the broader scientific community, which can better align research with issues of public importance.

While this step is presented last, it should occur throughout project development. Continuous communication brings scientific findings to the relevant audience and is especially powerful when data can be transferred in (quasi-)real time. This allows timely communication on data streams that may be of urgent need, such as the impact of extreme weather events, and can increase the likelihood that relevant actions are taken by stakeholders.

### Case Studies Implementing Workflow

#### Bold numbers in parentheses indicate the workflow step(s) being described

##### Belgium

**(1&2)** *CurieuzeNeuzen in de Tuin* (CNIDT, ‘*Nosy Parkers in the Garden*’), as described by Lembrechts et al. (2022), was a large-scale citizen science project designed to monitor the impact of extreme weather events on the microclimate of private and communal gardens in Flanders, Belgium (Figure 2; Table S1). The project served as a vehicle to raise societal awareness on the topic of climate change, and to investigate the potential of Flemish (peri-)urban nature (covering 12.5% of the region’s surface area) for climate adaptation and mitigation strategies. To be sufficiently representative of the variation across Flanders and, more importantly, involve a critical mass of the regions’ inhabitants, we established an IoT-enabled network across 5,000 locations. The IoT-connectivity enabled real-time data access, which, in turn, helped stimulate dialogue on localised weather impacts at the societal level. The communicative strength of the project was enhanced by prioritising communal gardens (e.g., parks, schools), and through publicly available interactive maps. Given the scale of the envisioned network, we made partnerships with academic institutions, civil society (citizen scientists, journalists), NGOs (e.g., Natuurpunt), government agencies (e.g., the Flemish Environmental Agency), and private partners (e.g., cell network provider Orange Belgium). Stakeholder meetings were organised regularly to communicate progress made and to exchange feedback.

The project comprised two measurement campaigns (April-October) in 2021 and 2022. **(3)** The key variables of interest were selected based on three categories: the degree of urbanity (e.g., imperviousness), the location’s geographical features (e.g., topography, geographical location), and other key drivers of microclimate (e.g., tree coverage, soil). This selection balanced high-resolution microclimate (meta)data for comparing local management practices with citizen science deployment (on their own properties) across the entire region. **(4)** Locations were selected based on remote sensing products and surveys completed by over 50,000 candidate participants upon registration. We used the code supplement here to distribute 5,000 sensors according to environmental variation (i.e., the first three components of a Factor Analysis of Mixed Data, Lê *et al*. 2008). At each selected location, a TMS-NB (TMS-4, Wild *et al*. 2019, with IoT connectivity) data logger was installed by a citizen scientist. Each participant paid €20 as a commitment fee, which could be reclaimed at the end of the project as a voucher, provided adequate completion of their measurements. **(5-7)** Preceding the measurement campaigns, a pilot study was conducted in and around the city of Leuven (Belgium) to finetune data logger installation for citizen scientists. Although this trial did not yet include IoT connectivity, it provided a crucial basis upon which the data management pipeline (e.g., data format, error flagging, dashboards) of the project was built. **(8-9)** Participants were prompted to complete two surveys, conducted in parallel with the microclimate measurements. Given the limitations of satellite-derived maps in capturing information on garden infrastructure and management, the engagement of citizen scientists was crucial for acquiring these data. During the measurement campaigns, each participant had access to their data via an online dashboard, which they could share freely. Including a national newspaper among the project’s stakeholders enabled regular updates of our measurements to the general public, highlighting weather extremes that occurred during the project. Between measurement campaigns and upon completion of the project, each participant was given a personalised report. We ended by organising a conference for partners and participants to present two years of microclimate citizen science, including testimonials and the main takeaways of the project.

##### Oman

**(1)** The second case study aimed to describe the microsites, distribution and climate sensitivity of plants in the Western Hajar Mountains of Oman. The landscape has complex geography with high mountain peaks (600 m-3000 m) divided by deep, seasonally dry riverbeds, with sparse and scattered grasses, shrubs and trees. The climate is extreme with summer temperatures reaching >50°C and occasional winter snow on the highest peaks, yet minimal rainfall of ∼225 mm of rain a year (World Bank Group 2024). Due to the harsh climate, complex geography and our pre-existing knowledge of the Hajar Mountains as a refugium, we expected microclimate to strongly influence the distribution of montane plants. We recorded vegetation in 100 m^2^ plots, alongside microclimate sensors measuring air temperature (TC Direct Ultra Fine Wire Type K Thermocouple) and relative humidity (Kestrel DROP D2). **(2)** From the outset we worked with the Oman Botanic Garden field botany team, who provided expert local knowledge, assistance with field surveys and steered research aims. They confirmed that the basic ecology and distribution of many of the plants in the Hajar is poorly known, which makes their conservation challenging. Therefore we aimed to measure the microclimatic niches these plants occupied, to guide *ex-situ* cultivation and construct species distribution models to better target surveys, designate protected areas and predict the effects of climate change. The Omani team highlighted the need to consider the role of humidity in shaping microclimate variation and species’ distributions, because occult precipitation (condensing water vapour) is the primary moisture source in coastal arid regions of Oman, and is not quantified in remote-sensed precipitation layers. Local insight was also crucial to avoiding military exclusion zones and ensuring safe import of sensors into the country.

**(3&4)** Plot sites were initially chosen based on a precursor to the code supplement, employing topographic, vegetation cover, macroclimate (temperature and humidity), and solar constant, as locally important microclimatic drivers. Plots were distributed randomly throughout the microclimate bands identified during the ordination process. We included redundant plots within each band, in anticipation that some plots would be inaccessible due to unmapped cliffs and ravines. **(5-7)** Data were collected in 160 plots using the mobile app ArcGIS Survey123, which allowed direct upload of GPS position and uploaded data to cloud storage, as well as ArcGIS Field Maps, which allowed survey maps to be checked easily in the field. Even with partnership knowledge and previous experience in the region, we found that travelling to most of the randomised plots was unfeasible. Hence, we used ArcGIS Field Maps to identify alternative accessible areas for each band, within which we aimed to measure at least 10 plots to ensure that plots were still representative of the full range of microclimatic variables across the landscape. **(8)** Some logger deployments were successful, however not all sensors were deployed due to logger battery damage during transit. **(9)** We presented preliminary findings in person to the Oman Botanic Garden, and have since summarised the results in a Master’s thesis shared with the Omani team. We are now in the process of co-designing a national Omani microclimate monitoring network with the Oman Botanic Garden.

##### Madagascar

**(1)** Our case study in southeastern Madagascar explored how microclimate drives reptile and amphibian (herp) diversity and distributions. The region experiences dramatic seasonal rain and inclement weather, including frequent cyclones (several during deployment). Prior to sensor deployment, we selected a 16-km^2^ study community, Ambalavero, based on its accessibility (a maintained hiking trail, as no roads exist in the region) and from observed environmental heterogeneity from prior visits. Ambalavero features rolling hills and cliffs with a mosaic of old-growth broadleaf rainforest, patches of selectively-logged native forest, smallholder agricultural plots (primarily rice paddy and livestock pasture), and “fallow” land of regrowing native and invasive vegetation. Given our focus on herp ecology, we monitored air temperature and relative humidity, both near-surface and within canopies for forested sites (at least 3m height). All navigation to and around the landscape was necessarily done by foot. **(2)** Early in project planning, we consulted the local community forest management organisation (“COBA”) to request permission to access community forests, discuss research priorities, and pay permit fees. As some forest patches in the region are considered sacred, it was paramount to identify where foreign researchers were allowed, and what rituals were necessary prior to entering forests. We allocated part of the project budget to compensation for landowners whose properties were chosen for sensor deployment. Project supervision, design, logistics, and implementation occurred over a two-year timeline (one five-month field campaign with several shorter field campaigns) to lay groundwork for long-term microclimate monitoring.

**(2)** We compiled gridded layers of environmental drivers of interest – elevation, slope, aspect, normalised difference vegetation index (NDVI), distance to nearest forest, and categorical land cover – that are known drivers of both microclimate and herp distributions (Campbell & Norman 2012; Nowakowski *et al*. 2017). **(4)** The exact budget allocated to sensors was flexible. As a result, we used the code supplement to designate the required number of sensors that adequately represented environmental variation. Through our algorithm, we designated 75 possible locations for sensors. **(5)** We performed *ex-situ* pilot studies to estimate battery duration and thermal accuracy of several microclimate sensor models, including previously purchased equipment, through which we chose Onset HOBO Pro and Pendant sensors supplemented by several Lascar EL-USB sensors with ultra fine-wire thermocouples. **(6&7)** When we began deployment, we deemed some sites to be ineligible or inaccessible, particularly those on cliff faces. After deploying sensors at eligible sites, we used the code supplement to iteratively select new sites that best complemented existing deployments (re-running code while camping in the field), and with updated criteria (e.g. a threshold of 45° slope to avoid cliffs). This resulted in a final set of 54 locations, 19 of which had multiple vertical strata (Figure 2). For sites located within community forests, coordinates and verbal directions were provided to the COBA president. For sites on private property, we greeted landowners (always with a team member who resided locally and was familiar with most families) and provided an introduction of our backgrounds, research aims, and methods. Such in-person consultations were necessary as there was no Internet or cell service to contact landowners prior to visit. We asked permission of landowners to deploy sensors, and offered monthly compensation. We also encouraged landowners to inspect sensors periodically, and to notify us of damage or theft. All participants responded enthusiastically and most indicated they were proud to take part in a community initiative that would encourage future research. **(8&9)** In two years of monitoring, two sensors were water damaged, and none have been lost or stolen. Microclimate monitoring continues at present, and we hold annual meetings with COBA members to provide updates on results and to reaffirm land access permission.

### Challenges of establishing and managing microclimate networks

When establishing and maintaining a microclimate network, various obstacles may arise that extend beyond any of the individual steps in the workflow described above. Here, we briefly discuss some challenges in deploying and operating microclimate networks: (i) the discrepancy between the resolution of remotely sensed and microclimate data, (ii) the trade-off between project budgeting and the network’s size, and (iii) the challenges of maintaining microclimate networks based on citizen participation.

(i) Remotely sensed gridded products are often used as predictors in spatial ecology, but are typically of far coarser resolution (1-100 km) than the scales at which microclimates are generated (1-100 m; Geiger 1942). Thus, the remote sensing layers provided as input to our code supplement may not be entirely reliable for capturing true landscape-scale environmental variation. While disaggregation and interpolation (e.g., bilinear, nearest neighbour) are computationally feasible, these may only be ecologically relevant if the input layers are already relatively fine-scaled and adequately capture the key drivers of microclimate (Klinges *et al*. 2024). A more representative solution could be to use gridded products with higher resolution to compute new (proxy) covariates. For instance, if precipitation or soil moisture data of an appropriate resolution are unavailable, high resolution Topographic Wetness Index (TWI, representing the likelihood of a given area to be wet; Beven & Kirkby 1979) may be calculated from topographic data (e.g. function ‘.topidx’ in *microclimf*, Maclean 2023). TWI as a fine-scale proxy for soil moisture would improve microclimate sensor placement compared to use of coarse soil-moisture or precipitation data. Once an initial microclimate network is in place, its measurements can also be leveraged for evaluating the explanatory power of remote sensing, to inform the next iteration of sensor deployments. Here, one can compare microclimate measurements from sensors deployed within similar conditions as quantified by spatial inputs (i.e. sensors placed within the same environmental ‘bin’ by the code supplement, Step 4, Figures S1-S2); similar microclimate measurements between these sensors would suggest current spatial inputs are satisfactory, while divergent measurements suggests important environmental variation has been left unexplained by current inputs. Ultimately, the importance of some microclimate drivers may remain partially unknown until sensors are deployed, again pointing to the value of pilot studies and iterative deployments.

(ii) A typical challenge lies in balancing resources within a project’s budget. Given the auxiliary expenses that may arise beyond sensor purchases – including compensating stakeholders and landowners for their invested time when appropriate, as we advocate for – we encourage researchers to keep project scopes modest for a first sensor implementation, especially concerning the spatial extent. While some researchers may be tempted to cover a large area, including moving individual sensors across multiple locations, we encourage in most cases continuous measurements at fewer sites to prioritise quality over quantity. Sampling realistically-sized landscapes also facilitates sensor revisits, which is typically needed to monitor sensor (mal)function and for data downloading (but see “IoT as the Future of Sensor Networks” below). Maintaining realism early in study design provides greater flexibility in the later stages that we describe, therefore enabling data re-use and upwards scaling.

(iii) In recent years, citizen science has received growing recognition for large-scale data collection and science communication (Dickinson *et al*. 2010), but this comes at the cost of increased complexity. Measurement devices need to be distributed among participating citizens, who need to be trained to collect the required measurements. To further ensure the network’s longevity, citizen scientists need to remain committed, achieved through frequent rounds of stakeholder communication and feedback. Even if participants are properly trained, citizen-obtained data may be more prone to measurement errors, demanding a strong quality assurance workflow.

### Co-development across the Workflow with Stakeholders

Given that microclimate by definition changes across local spatial scales, access to and intimate knowledge of landscapes is important for understanding microclimate variation. Whether through working with land management organisations, local governments, Indigenous communities, or other stakeholders, grounding microclimate research in local understanding better ensures its continued relevance and impact. Drawing from other disciplines such as sociology and implementation science, ecologists measuring microclimate can meaningfully engage with communities when establishing sensor networks (Shinde *et al*. 2023).

Landowners or recreationalists familiar with a target landscape can provide suggestions on safe, accessible and interesting deployment locations. Informing community members early can also reduce theft or inform researchers of possible causes of incidental damage, such as trampling by livestock, areas that flood, or planned management or land development. Such interactions can also keep external researchers informed; in our Madagascar case study, every sensor had a local community member who was provided with a stipend to check regularly for sensor damage or failure. Sustaining such relationships can also broaden the impact of research through public education. Our Belgian case study developed multimedia to inform Belgian residents of the project, and the value of microclimate data for understanding urban heat waves and drought.

Co-development with stakeholders can also address power imbalances that have prevented local and global communities from fully benefiting from research findings. When external researchers do not consult local communities and scientists, particularly those of marginalised identities, environmental data collection emulates earlier extractions of raw materials (Baker *et al*. 2019). Community members who aid in environmental research but do not see its outcome can feel exploited (Dogan & Wood 2023). Frameworks to safeguard community data, such as the CARE principles for Indigenous data governance (Carroll *et al*. 2020), aim to prevent these and other extractive practices, for example by promoting the collective benefit from gathered data and establishing early on who has the authority to control the data. By integrating local community members into all stages of environmental sensor study design, researchers avoid blindspots, achieve more equitable outcomes, and can generate more impactful research (Reid *et al*. 2021).

### IoT as the Future of Sensor Networks

The methodology we provide here remains useful as sensor technologies advance in the future, such as with the increasing adoption of Internet of Things (IoT). Importantly, IoT-connected sensors would in many ways facilitate the implementation and especially maintenance of sensor networks.

IoT-connected sensor networks enable continuous sensor performance monitoring, reducing the number of field visits and minimising data loss from malfunctions. Moreover, real-time data transfer accelerates data analysis, shifting the focus from microclimate (long-term) to microweather (short-term) monitoring. Timely data access is invaluable for understanding extreme weather events’ impact on ecology and hastens communication with stakeholders. This rapid information dissemination enhances public understanding, informs decision-making, and expedites necessary management interventions – critical imperatives in an era marked by unprecedented climate change. IoT capabilities also expand deployment strategies, such as animal-mounted sensors to better capture proximal conditions (Ellis-Soto *et al*. 2023). Additionally, IoT sensing can provide metadata, such as battery level and network connectivity, to minimise wasted time in the field and open up opportunities for adaptive sensing. Such additional metadata facilitates improvements to sensor design and intercomparison of methods and findings across networks. Upscaled insights can then guide the establishment of regional microclimate networks as part of global microclimate measurement campaigns.

Pioneering initiatives with IoT-sensors, like the Flemish case study mentioned above, are already underway. However, while manuals for developing IoT sensors are available (e.g., Mickley *et al*. 2019), ecologists still lack commercially available, low-cost sensors with IoT-connection. This advancement is a crucial step toward creating a comprehensive network of ’microweather stations’ worldwide, much like the standardised weather station network currently in place but, ideally, with locations informed by environmental variables rather than administrative boundaries and socioeconomics (Lembrechts *et al*. 2021). Such developments are vital for evaluating microclimate’s role in ecological processes amid a changing climate, and integrating local measurements into global intercomparisons.

### Applying the Workflow to Other Environmental Sensing

While our focus has been on microclimate monitoring, the principles and solutions discussed here are relevant across a spectrum of environmental variables that vary significantly across small spatial extents, from air and water quality to noise pollution ). Implementing networks for other variables would likewise entail many of the approaches we highlight above.

In any case it is critical to identify the key drivers (natural and social) of spatiotemporal variation in the environmental variable of interest. For instance, creating a monitoring network for chemical pollution necessitates consideration of the specific chemical types, spatial and temporal distribution of pollutant sources, and the environmental spread rate (Artiola & Brusseau 2019). Our site selection, which currently relies heavily on remote sensing data, might not be sufficiently accurate to capture such variables in detail, and local knowledge may need to inform the spatiotemporal configuration of the monitoring campaign. Similarly, when addressing noise pollution and its effects on human well-being, subjective experiences play a pivotal role, often diverging from objective decibel measurements (Murphy & King 2022). In such scenarios, a deeper understanding of human stakeholders’ perceptions becomes essential to align monitored sound levels with their lived reality. Despite these potential variations, our methodical approach – engaging stakeholders, identifying driving factors, and leveraging them for selecting measurement locations – promises to be advantageous across almost all environmental applications.

### Synthesis and Conclusion

The increased availability of small, low-cost and wireless environmental sensors has revolutionised ecological research and conservation efforts, enabling autonomous data collection across many study sites. Despite this advancement, ad-hoc approaches to selecting deployment sites and measurement methodologies have constrained the full potential of sensor data in applied ecological contexts. While ample resources and robust methodologies exist for guiding site selection from the perspective of biodiversity surveys (e.g., Carvalho *et al*. 2016), there is a notable scarcity of guidelines specifically tailored to inform the strategic placement of sensors to monitor key environmental variables (Lembrechts *et al*. 2021).

Our paper addresses this gap by providing a practical workflow for microclimate sensor networks from conception to execution to dissemination, promoting improved standardisation in environmental monitoring across studies. Crucially, our workflow emphasises effective communication and collaboration with stakeholders, fostering ownership and support among researchers, policymakers, and community members. This collaborative approach is vital for translating sensor data into actionable, applied insights that inform evidence-based policies and practices for managing anthropogenic stressors, such as habitat loss, ecosystem degradation, climate change, and overexploitation. By following our systematic approach, researchers can ensure that sensor deployment is well-informed, scientifically robust, and aligned with project and stakeholder goals and budget constraints, ultimately enhancing the quality, utility and longevity of environmental sensor data with local practitioners in mind.

## Funding

DHK was supported by The Explorer’s Club Fjällräven Field Grant, The University of Florida TCD Field Research Grant, and the National Science Foundation (DGE-1842473). JJL and SVdV were supported by the Research Foundation Flanders (W001919N, G018919N and 1S90923N), and the BiodivERsA-project Forest-Web-3.0 (G0GDZ23N). KH-C and RS were supported by the Royal Geographical Society (with IBG) Small Research Grant, and by The British Omani Society (formerly The Anglo-Omani Society).

## Author Contributions

DHK, JJL and RS conceived the study. DHK, JJL, SVdV, KH-C, and RS conducted fieldwork. DHK led software development with input from all authors, especially SVdV. All authors contributed substantially to manuscript original drafting and revision. RS supervised this work and should be considered senior author. Our study brings together multinational authors, and is informed by collaborative ongoing work with researchers local to the case study regions provided here.

## Data and Software Availability

No new data were generated as part of this manuscript. Software for implementing our workflow is provided via a GitHub repository at https://github.com/dklinges9/Microclimate-Sensor-Networks.

## Supplementary Information

**Table S1.** Summary information for each case study region for which the workflow was implemented.

| Location | Area (km<sup>2</sup>) | Number of plots | Duration | Sampling rate | Ecological processes of interest | Measurements |
| --- | --- | --- | --- | --- | --- | --- |
| Flanders, Belgium | 13,620 | 5000 | 6 months x2 | 15 mins | Ecosystem functioning of gardens | Air temperature (15cm aboveground), soil moisture, composite soil sample, soil temperature (2-15cm belowground) |
| Madagascar | 16 | 72 | 3 years | 15 mins | Amphibian distributions and diversity; agricultural productivity; climate refugia of sacred forests | Air temperature and relative humidity (15cm aboveground), soil temperature (5cm belowground) |
| Oman | 190 | 160 | 1 month | 17 mins | Plant species distributions and plant diversity distributions, Identify future climatic refugia for montane plant species | Relative humidity, air temperature (20cm aboveground) |

**Figure S1.**
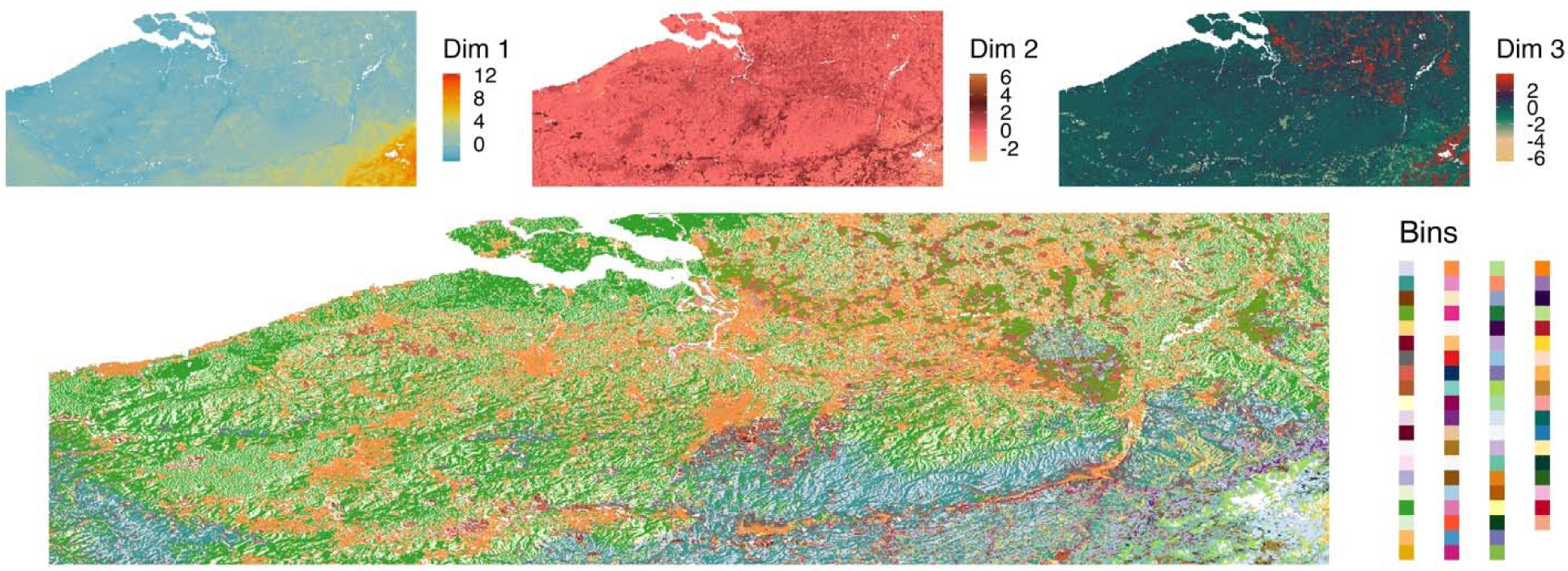
Example visualisations that are auto-generated from the code supplement, here for Flanders, Belgium (50° 46’ 3.28175" S, 51° 30’ 20.88533" S, 2° 31’ 32.46287" E, 5° 58’ 14.61288" E). Displayed are the values for each of the three dimensions as part of the ordination (top row), as well as the bin identity for each spatial pixel (bottom row, 78 bins here). Sensor sites are then selected to equally represent environment space across these bins (e.g. one sensor per bin).

**Figure S2.**
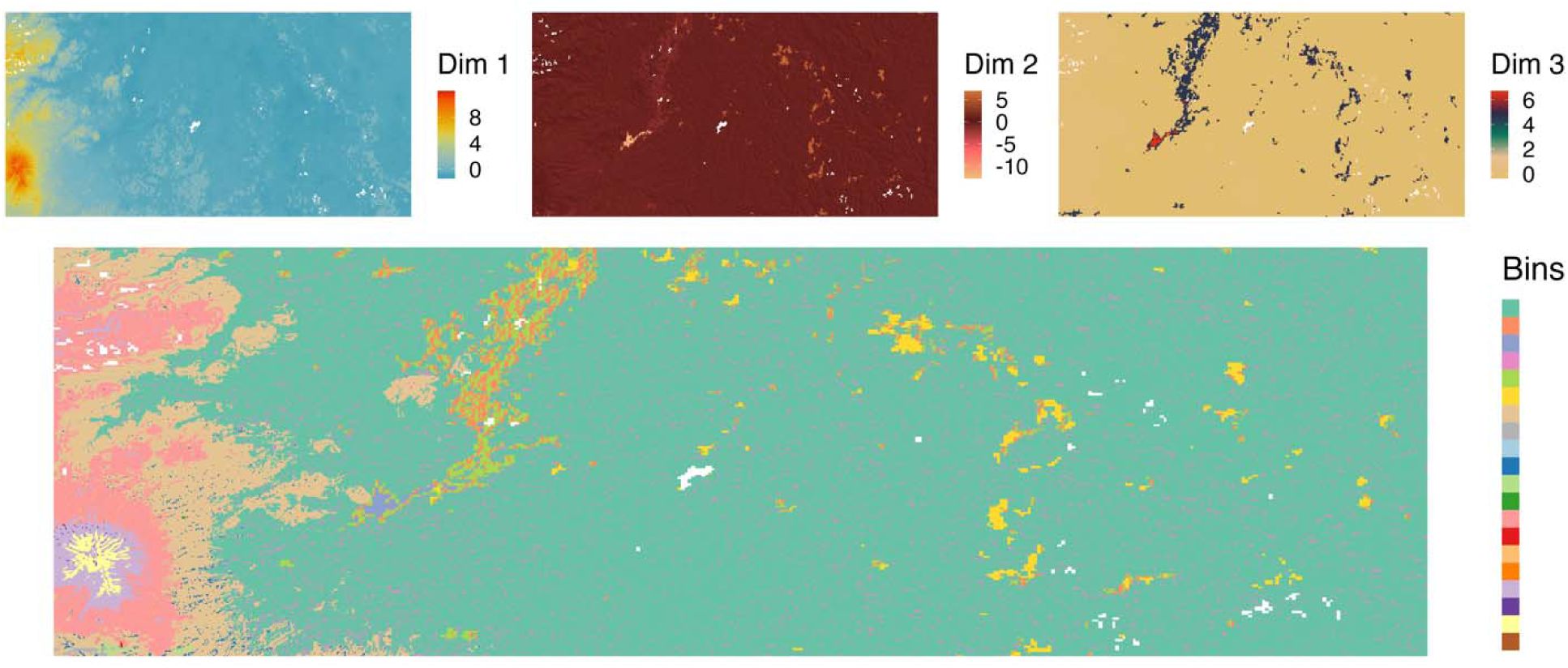
Example visualisations that are auto-generated from the code supplement, here for South Sumatra, Indonesia (-3° 26’ 25.368" S, -3° 0’ 0" S, 102° 34’ 58.8" E, 103° 43’ 40.8" E). Displayed are the values for each of the three dimensions as part of the ordination (top row), as well as the bin identity for each spatial pixel (bottom row, 20 bins here). Sensor sites are then selected to equally represent environment space across these bins (e.g. one sensor per bin).

**Figure S3.**
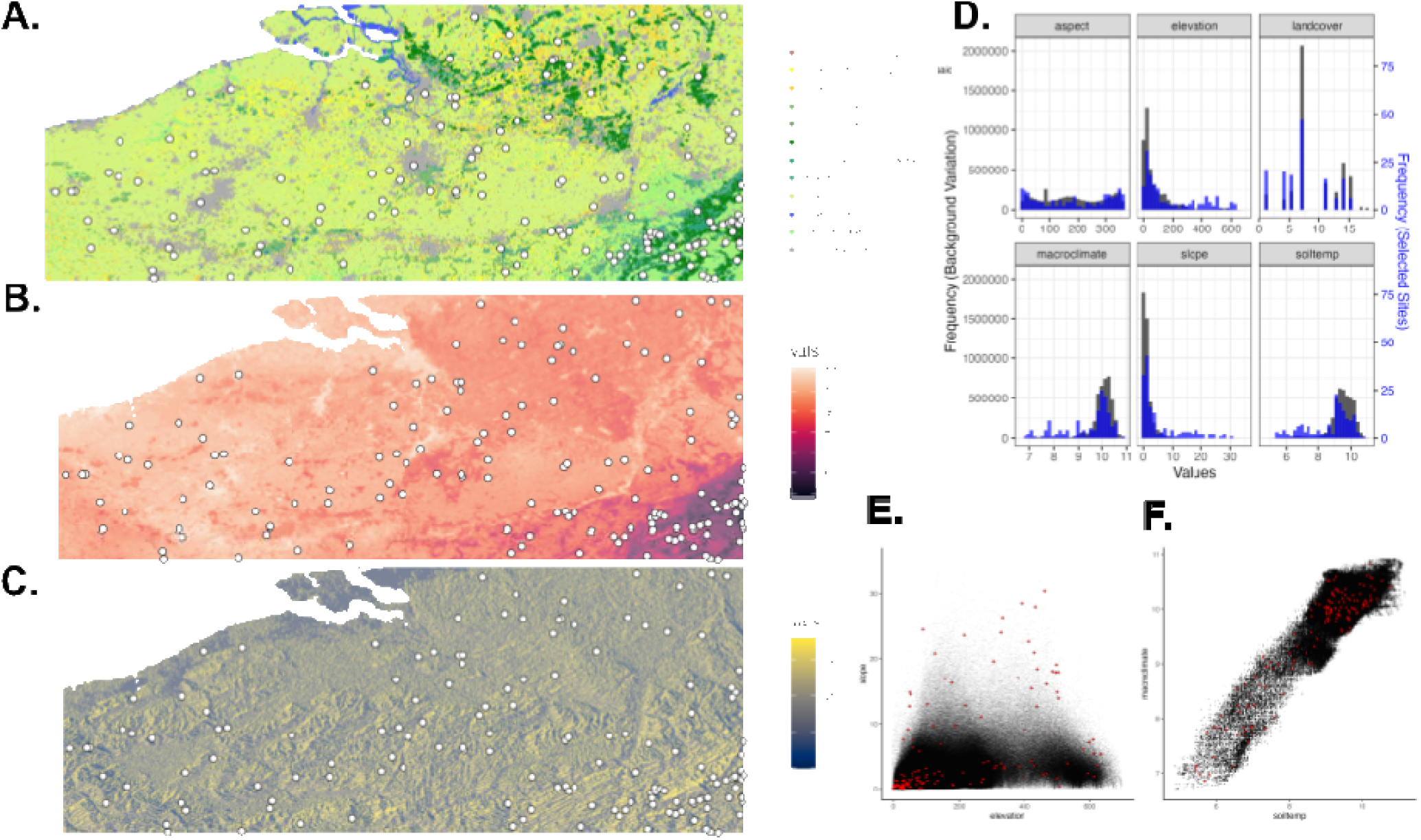
Example visualisations that are auto-generated from the code supplement, here for a sensor site selection (points on maps) conducted in Flanders, Belgium. A: categorical land cover as estimated by Copernicus CCI-LC; B: average soil temperatures (Lembrechts et al. 2022); C: aspect estimated from a 100-m DEM derived from Amazon Web Services; D: histograms depicting the environmental representation of sites chosen for sensors (blue bars) with background environmental variation (grey bars); scatter plots of chosen sites (red points) and background variation (black points), here depicted for relationships of elevation with slope (E) and WorldClim macroclimate with soil temperature (F).

**Figure S4.**
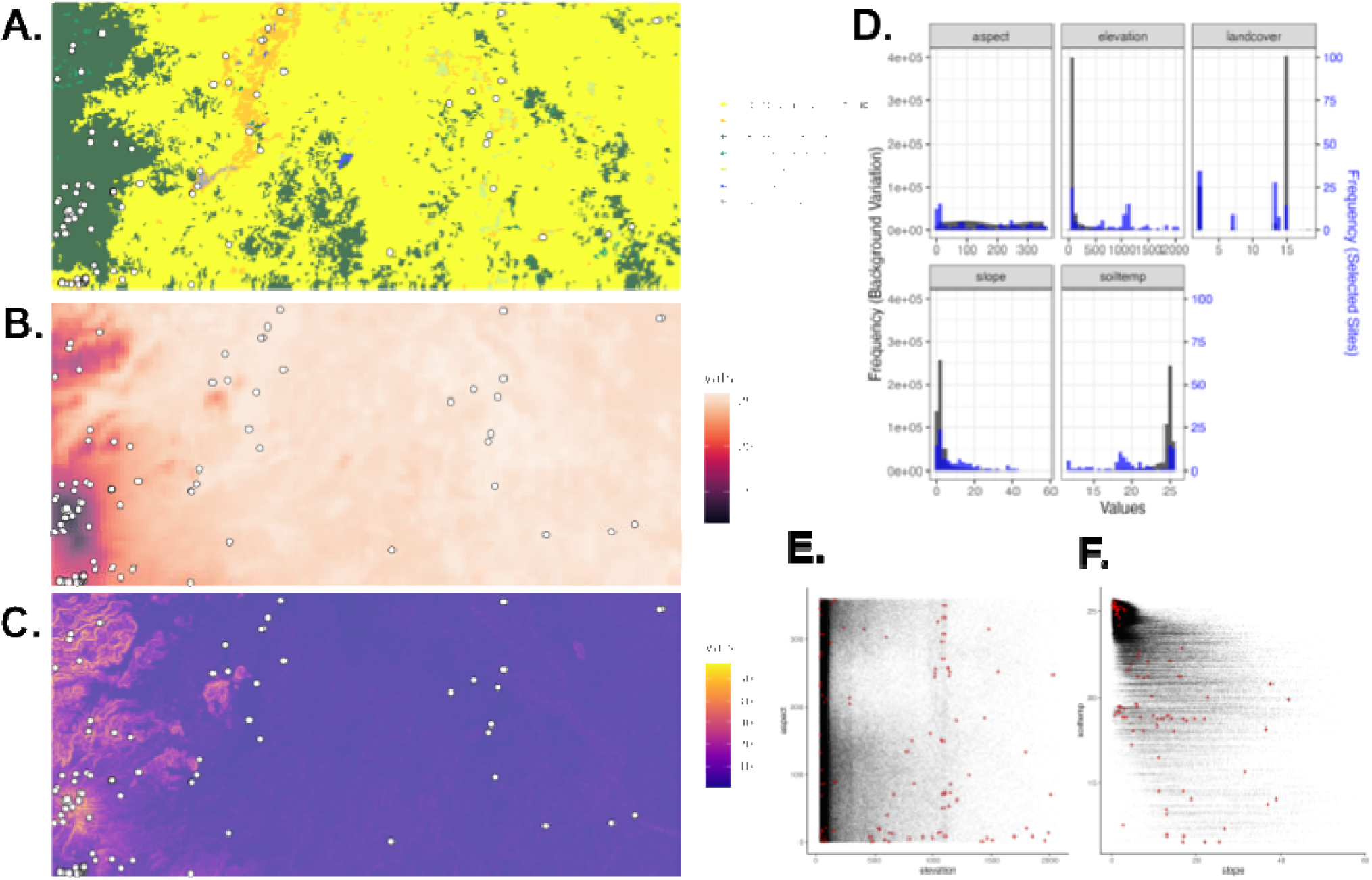
Example visualisations that are auto-generated from the code supplement, here for site selection for 93 sensor (points on maps) conducted in South Sumatra, Indonesia (-3° 26’ 25.368" S, -3° 0’ 0" S, 102° 34’ 58.8" E, 103° 43’ 40.8" E). A: categorical land cover as estimated by Copernicus CCI-LC; B: average soil temperatures (Lembrechts et al. 2022); C: slopes estimated from a 100-m DEM derived from Amazon Web Services; D: histograms depicting the environmental representation of sites chosen for sensors (blue bars) with background environmental variation (grey bars); scatter plots of chosen sites (red points) and background variation (black points) here depicted for relationships of aspect with elevation (E) and slope with soil temperature (F).

## Notes

https://github.com/dklinges9/Microclimate-Sensor-Networks

### Competing Interest Statement

The authors have declared no competing interest.

